# Physics-Informed Estimation of Electrostatic Attraction During Fingertip Sliding Under Varying Speed and Normal Force

**DOI:** 10.64898/2026.08.05.743019

**Authors:** Celal Umut Kenanoglu, Yasemin Vardar

**Affiliations:** Department of Cognitive Robotics, Delft University of Technology, Mekelweg 5, Delft, 2628 CD, The Netherlands

**Author notes:** Corresponding author (Y. Vardar).

**Keywords:** Electrostatic actuation, electroadhesion, contact mechanics, fingertip-surface interaction

## Abstract

Electrostatic actuation is an emerging technology for generating tactile sensations on capacitive touchscreens through voltage-induced attractive forces between a fingertip and the surface. However, accurate control of electrostatic attraction during natural touchscreen interactions remains challenging because the applied normal force and sliding speed continuously vary, and their effects on the fingertip–screen contact and resulting actuation strength are not fully characterized. Here, we show how normal force and sliding speed systematically alter fingertip– screen contact area and electrical impedance, and use these measured changes to estimate electrostatic attraction during sliding. Contact area, interaction forces, and electrical impedance were measured simultaneously as participants slid their fingertips across an electrostatic surface under systematically varied normal forces and sliding speeds. These measurements revealed condition-dependent changes in fingertip contact, electrical interaction impedance, effective capacitance, derived effective gap thickness, and electrostatic attraction. We then incorporated these measured contact quantities into a physics-informed, data-driven model based on parallel-plate capacitor theory, in which effective capacitance, apparent contact area, and effective voltage determine the estimated electrostatic attraction. The resulting model links force- and speed-dependent changes in these quantities to electrostatic attraction while accounting for inter-participant variability through a participant-specific scaling factor. These findings provide experimentally grounded guidance for designing electrostatic surface-haptic feedback and future adaptive control strategies under realistic touch conditions.

## 1. Introduction

Modern interactive devices can deliver rich audiovisual content, yet their tactile output is often limited to simple vibrating alerts, far from the nuanced cues experienced when interacting with real surfaces. Electrostatic actuation offers a promising approach for rendering such cues by generating controllable forces at the fingertip-surface contact. This technology is particularly compatible with surfaces that consist of a conductive electrode covered by an insulating layer (e.g., touchscreens), where applying voltage produces an attractive electrostatic force during fingertip contact [1, 2]. The resulting attraction is governed not only by the applied voltage but also by the mechanical and electrical quantities of the fingertip–surface contact during sliding, including the contact area of the fingertip and the electrical impedance. As the fingertip is compliant and viscoelastic, these quantities vary with applied normal force and sliding speed [3, 4, 5, 6, 7, 8], making electrostatic attraction difficult to estimate and control under realistic touch interactions.

While prior studies demonstrated that applied normal force and sliding speed affect electrostatic actuation, the evidence remains scattered across separate measurements of contact area, electrical impedance, and electrostatic attraction force, with limited models connecting these quantities under the same sliding conditions. Increasing applied normal force has been generally associated with stronger electrostatic attraction (often inferred from tangential-force changes) [9, 10]. This trend was shown to persist even when the apparent contact area (i.e., fingertip boundary) was held constant [11, 12]. Sliding speed, in turn, has been found to reshape the dynamics of electrovibration-induced friction by shifting the interaction cutoff frequency, whereas normal force had a limited influence on these dynamics within the tested ranges [13]. Complementary electrical characterizations showed that lateral motion substantially increases the overall electrical impedance during sliding relative to stationary contact, while higher normal force can reduce impedance under moving conditions [14, 15].

These findings indicate that the intensity of electrostatic attraction depends strongly on variations in contact area and electrical impedance, which result from changes in normal force and sliding speed. However, how these contact quantities vary together across combined force–speed conditions, and how these variations can be parameterized for estimating electrostatic attraction during sliding, remains insufficiently characterized. A compact force- and speed-dependent model based on these contact area and electrical impedance is therefore still lacking. Existing modeling approaches have typically interpreted electrostatic actuation primarily through friction measurements, using either multiscale contact mechanics models [16, 17, 18] or parallel-plate capacitor theory of the fingertip–surface contact [19, 20, 21, 22, 23].

In the multiscale contact models, the real contact area is described across multiple length scales using fingertip properties and surface roughness. The applied voltage is then introduced as an additional electrostatic attraction that increases the real contact area, thereby altering friction [16, 17]. These models provide important physical insight into electroadhesive contact, but they require participant- and surface-specific parameters, such as skin mechanical properties, surface roughness, and contact geometry. These parameters are difficult to obtain routinely during dynamic touch and may vary across users and interaction conditions. Therefore, using such models for force estimation or adaptive control across changing force–speed conditions is not straightforward. In addition, studies using this approach have rarely combined simultaneous measurements of contact area and friction under systematically varied normal force and sliding speed conditions, and have typically been validated with a limited number of participants.

Alternatively, parallel-plate capacitor theory represents the fingertip–surface contact as two separated conductive plates. In this model, electrostatic attraction primarily depends on the applied voltage, effective contact area, gap thickness between the conductive layer of the fingertip and the surface, and the dielectric properties of the insulating layers, namely the touchscreen insulator and interfacial gap [10, 23]. Existing implementations, however, commonly rely on idealized assumptions, such as fixed effective contact area and gap thickness, despite well-known load- and motion-dependent changes in contact formation [4, 5] and electrostatic-actuation-induced changes in contact [24]. Consequently, limited knowledge of these evolving contact quantities limits the ability of these models to estimate electrostatic attraction under varying sliding conditions. Combined measurements of electrical impedance, contact area, and interaction force can provide the condition-dependent quantities needed for such estimation.

Addressing this limitation requires simultaneous measurements of contact area, electrical impedance, and interaction force under controlled force–speed conditions. Such measurements are needed to determine how normal force and sliding speed alter fingertip–surface contact area and electrical impedance, and to parameterize a compact model for estimating electrostatic attraction. The need for such a model is further amplified by well-known inter-participant variability in fingertip–surface interaction and electrostatic actuation, which calls for approaches that remain compact yet adaptable across users [13, 15, 24].

Here, we present a physics-informed, data-driven model of electrostatic attraction during sliding that integrates simultaneous measurements of the fingertip–screen contact under systematically varied normal force and sliding speed. We first quantify how key contact quantities (contact area, tangential force, electrostatic attraction, electrical impedance, effective capacitance, and effective gap thickness) change with normal force and sliding speed. We then incorporate these quantities into a parallel-plate capacitance framework, in which normal force and sliding speed affect the estimated electrostatic attraction via condition-dependent changes in effective capacitance, apparent contact area, and effective voltage. A participant-specific scaling factor accounts for differences between users while preserving the same force–speed relation across participants. Overall, this study demonstrates how interaction conditions reshape the fingertip–screen contact and provides a basis for adaptive control strategies that adjust electrostatic actuation based on force, speed, and user variability during realistic touch interactions.

## 2. Materials and Methods

### 2.1. Experimental Setup

During the experiments, the same experimental setup introduced in [24] was used (Fig. 1 and Fig. S1). Participants slid their right index finger across a capacitive touchscreen (SCT3250, 3M). The touchscreen was excited by applying an alternating voltage to its conductive layer. The excitation signal was generated in MATLAB/Simulink and output through a connector block (SCB-68A, NI Inc.) at a sampling rate of 10 kHz before being amplified with a high-voltage amplifier (9200A, Tabor Electronics). Participants wore a grounding wrist strap with an internal electrical resistance of 1 MΩ. The touchscreen was rigidly mounted onto two six-axis force/torque sensors (Nano17 Titanium, ATI Industrial Automation). The interaction forces were recorded using a separate data-acquisition card (PCIe-6321, NI Inc.) at 10 kHz. The participants’ finger motion was restricted to an approach angle of 60° and imposed by a motorized linear stage (NRT150/M, Thorlabs). The current was estimated from the voltage drop across a 10 kΩ series shunt resistor placed between the amplifier output and the touchscreen. The shunt voltage was measured using a differential probe (TA043, Pico Technology). Contact area was recorded from below using FTIR imaging [25] with a high-speed camera (MotionBLITZ EoSens mini2, Mikrotron) and lens (LM16HC, Kowa) at 1000 frames per second (fps). The glass plate was illuminated by a collimated LED light source (KL 2500, Schott) coupled with a diffuser. The entire assembly was mounted on damped posts on an optical breadboard to reduce the effects of external vibrations.

**Figure 1:**
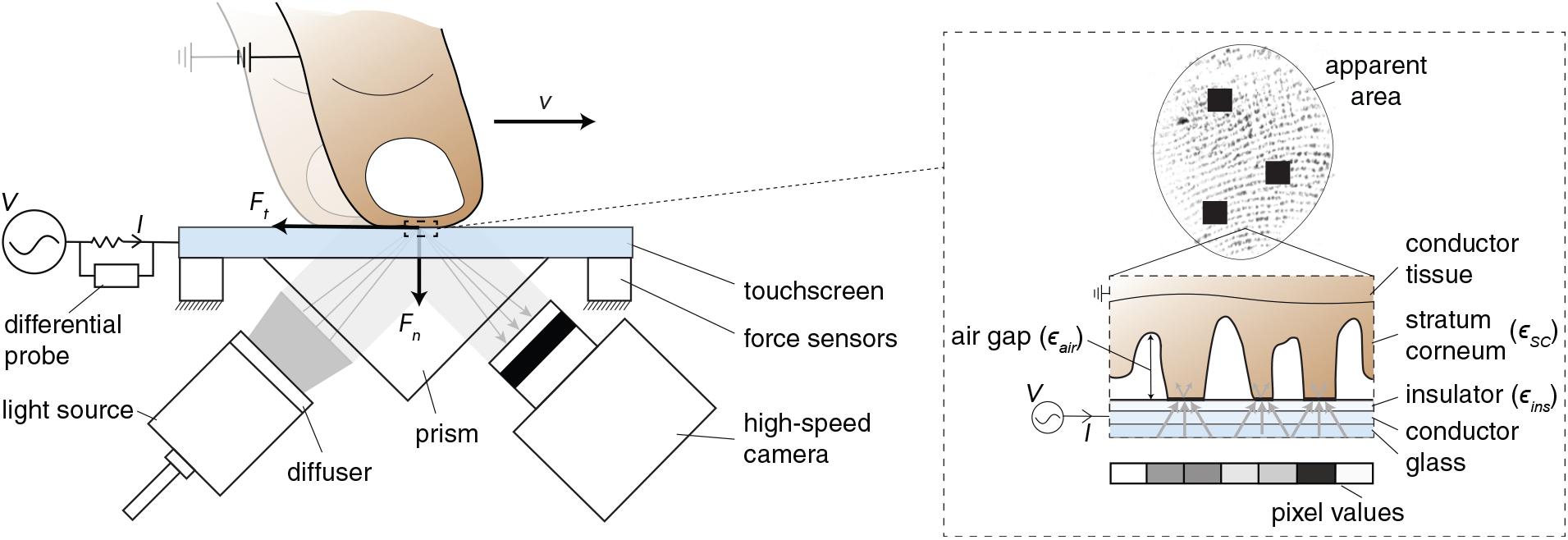
Illustration of the experimental setup. A fingertip slides on a capacitive touchscreen driven by an input AC voltage, while normal (*F*_*n*_) and tangential (*F*_*t*_) forces are measured with two force sensors. The normal force and sliding speed (*v*) are systematically varied. The input voltage (*V*) and resulting current (*I*) are recorded to quantify the electrical impedance. Finger–glass contact is imaged through a prism using frustrated total internal reflection (FTIR) with diffused illumination and a high-speed camera [25], where dark pixels indicate intimate contact (optically-resolved real area) and the fingertip boundary defines the apparent contact area. The black squares on the image are deliberately added to protect the sensitive fingerprint data. *ε*_ins_, *ε*_SC_, and *ε*_air_ denote the relative permittivities of the insulating layer, stratum corneum, and air gap, respectively.

### 2.2. Participants

The experiment was conducted with seven men and three women with a mean age of 27, SD:±2.45. The study was conducted in accordance with the Declaration of Helsinki and was approved by the Ethics Council of TU Delft (application no. 5108). All participants provided informed consent.

One participant (male) was excluded from the main quantitative analyses due to the pronounced moisture accumulation observed at the fingertip–surface contact during the experiment and across all experimental conditions. This moisture accumulation was clearly visible in the FTIR images as condensation-like regions beneath the fingerprint. The same participant also showed a markedly different electrical response, with substantially lower interaction impedance and higher effective capacitance (approximately 4–10-fold differences relative to the remaining cohort), and the electrostatic attraction response was close to zero in several force–speed conditions. These observations indicate that the intended dry-contact electrostatic actuation condition was not satisfied [24, 26]. Therefore, all main results are reported for the remaining nine participants.

### 2.3. Procedure

Before experiments, participants washed their hands and dried their fingertips with a microfiber cloth. In each trial, the finger was moved laterally at a constant speed (10, 20, or 30 mm/s) using the motorized linear stage. During this motion, participants regulated their normal forces to a target value (0.5, 1.0, or 1.5 N) using real-time LED feedback computed from the force-sensor signal. Yellow indicated forces more than 10% below the target, green indicated forces within the ±10% tolerance band, and red indicated forces above the target. Each force–speed condition was repeated three times. Results were collected only if the measured normal force varied by no more than 10% from the target and the fingerprint was sufficiently visible for contact area estimation. Within the same sliding stroke, electrostatic actuation was cycled between voltage-off and voltage-on conditions (100 V_peak_, 75 Hz sine wave). The stimulus frequency was selected based on setup characterization that showed a relatively stable response in the corresponding force-frequency range, thereby reducing the influence of setup dynamics on the measured signal [24] (Fig. S2). Moreover, frequency-dependent impedance measurements conducted in an earlier study [24] also showed that the effective capacitance exhibited its strongest variation at lower frequencies and became comparatively less frequency-sensitive above approximately 70 Hz (Fig. S3). The selected frequency and amplitude also lie within a perceptually relevant range for finger-electrostatic surface interaction [27]. To reduce moisture-related variability across trials, a fan was used to help maintain consistent skin dryness. Each participant completed the full experimental session in about 30 minutes.

### 2.4. Data Analysis

Data from the force sensors, motion-stage commands, and camera triggering were synchronized through MATLAB/Simulink. Signals other than the contact images were both generated and logged in Simulink, while image acquisition was handled using MotionBLITZ software. A trigger signal was sent from Simulink once the finger reached the camera’s field of view, and only the corresponding segments of the normal (*F*_*n*_) and tangential (*F*_*t*_) force measurements were retained for subsequent analysis. More details of signal processing, data extraction, and analysis are provided in the Supplementary Text (Figs. S4–S6).

Prior to contact-area estimation, the raw FTIR images were geometrically rectified. Radial distortions caused by the camera lens were compensated using intrinsic calibration parameters obtained from a checkerboard-based calibration procedure. Subsequently, a projective transformation was applied to generate an equivalent top-view representation [28]. The homography matrix was determined by imaging a circular rubber marker placed on the glass surface and mapping its observed elliptical projection back to a circle.

The apparent contact area, *A*_app_, refers to the macroscopic contact region at the interface (i.e., the fingerpad boundary), whereas the real contact area corresponds to the aggregate of microscopic asperity contacts within this region. In this study, FTIR imaging was used to estimate the optically resolved contact area associated with intimate contact at the interface. Real contact area was computed from the FTIR images following the procedure in [29]. As the real contact area is inherently scale-dependent [30, 31], the optically resolved measured real contact area value in this paper should be regarded as an optical approximation of the true contact area. Therefore, throughout this paper, this optically resolved contact area is referred to as the measured real contact area.

For each experimental trial, the ratio of measured real contact area between voltage-on and voltage-off conditions was calculated as *A*^on^/*A*^off^, where *A*^on^ and *A*^off^ represent the mean contact areas measured during the corresponding phases of the same sliding motion. In addition, the electrostatic attraction force was estimated according to *F*_*e*_ = (1 −*µ*_off_ /*µ*_on_)*F*_*n*_ [10, 18, 24, 32], where the friction coefficient is defined as *µ* = *F*_*t*_/*F*_*n*_. Electrostatic pressure, defined as attraction per unit measured real contact area, was computed as *p*_*e*_ = *F*_*e*_/*A*^on^.

The voltage, *V*, and current, *I*, signals were analyzed at the electrostatic actuation frequency (*f*_0_ = 75 Hz). After removing the DC component, the complex voltage and current components at *f*_0_ were obtained from the Fourier transform of each voltage-on segment. The current was estimated from the voltage drop across the 10 kΩ shunt resistor, measured using a differential probe. The probe input impedance (4 MΩ *∥* 7 pF per input to ground) was much larger than the shunt resistance. Using the conservative input-resistance value of 4 MΩ, the corresponding probe-loading current was estimated to be below 0.25% of the shunt current and was therefore neglected.

The total complex impedance, *Z*_total_, was then calculated as

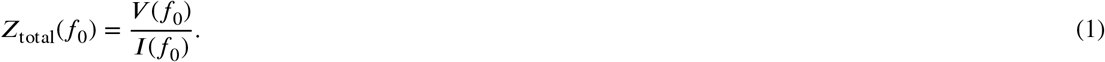

The known environmental impedance was removed from the measured total impedance:

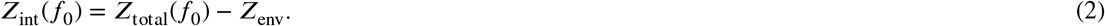

Here, *Z*_env_ accounts for the known dominant series resistance contribution of the grounding wristband and the shunt resistor used for current measurement. In this setup, *Z*_env_ was approximated by the known series resistances, *Z*_env_ ≈ *R*_wrist_ + *R*_shunt_ ≈ 1 MΩ + 10 kΩ. Thus, *Z*_int_ represents the impedance of the finger–touchscreen interaction after removing the known environmental contribution.

The measured interaction impedance was represented using an effective parallel RC model to obtain *C*_eff_ and *R*_eff_ (Fig. 2). The interaction impedance was converted to an admittance, *Y*_int_ :

**Figure 2:**
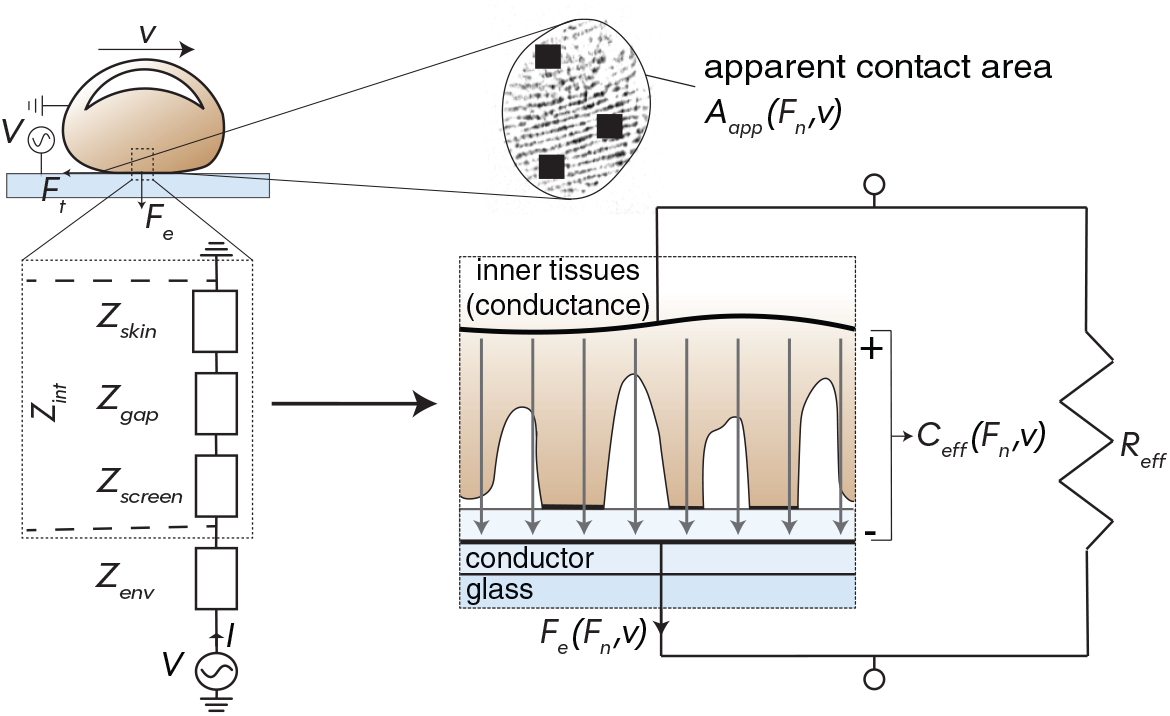
Physics-informed, data-driven model of electrostatic attraction during fingertip sliding. The fingertip–screen contact was represented by an electrical impedance network including skin (stratum corneum) *Z*_skin_, gap *Z*_gap_, insulator layer of screen *Z*_screen_, and environmental contributions *Z*_env_. The interaction impedance was then expressed as an effective parallel RC circuit, yielding the effective capacitance *C*_eff_ and effective resistance *R*_eff_. The effective capacitance was combined with the apparent contact area *A*_app_ and the voltage acting across the fingertip–screen contact in a parallel-plate capacitor framework to estimate the electrostatic attraction force, *F*_*e*_.

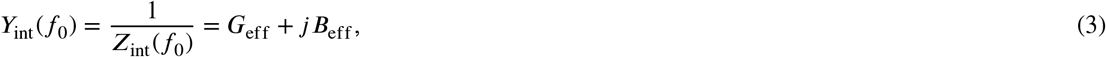

where *G*_eff_ and *B*_eff_ are the effective conductance and susceptance (imaginary admittance), respectively. The corresponding effective resistance and effective capacitance were defined as *R*_eff_ = 1/*G*_eff_ and *C*_eff_ = *B*_eff_ / (2π*f*_0_). This parallel R–C representation was used as an effective decomposition of the interaction admittance into conductive and quadrature components. The effective capacitance (*C*_eff_) was obtained from the quadrature component of the interaction admittance and should therefore be interpreted as a frequency-specific descriptor of the capacitive response of the fingertip–screen contact at the stimulation frequency, rather than as a direct measure of the physical finger–screen capacitance. In the literature, equivalent-circuit models sometimes include a small series resistance in addition to the parallel RC branch [15]. In the present analysis, this series resistance was neglected and *Z*_int_ was represented using an effective parallel RC model. This approximation is supported by previous equivalent-circuit fits showing that the fitted series resistance is only about 1% of the parallel-branch impedance under similar touch conditions [15]. A similar simplification has also been used in prior impedance measurements, where the body/internal-pathway impedance was considered negligible compared with the dominant interfacial impedance [33].

### 2.5. Modeling

#### Parallel-plate capacitor theory

The electrostatic attraction force is determined by the applied voltage, the effective contact area, the gap thickness between the dielectric layers, and the dielectric characteristics of the insulating layers between the conductive bodies. Based on parallel-plate capacitance theory, the electrostatic attraction force at the fingertip–screen contact can be calculated as follows [10, 21, 23]:

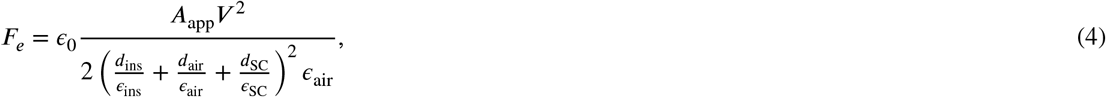

where *A*_app_ is the apparent contact area, which is considered as the macroscopic effective area participating in the electrostatic interaction, following previous studies that used parallel-plate formulations [10, 21, 23]. *V* is the effective voltage at the fingertip–screen contact, *d*_ins_, *d*_air_, and *d*_SC_ are the thicknesses of the touchscreen insulating layer, the air gap, and the stratum corneum, respectively; and *ε*_ins_, *ε*_air_, and *ε*_SC_ are the corresponding relative permittivities. Under the ideal assumption of negligible leakage, the capacitance of the same layered stack is

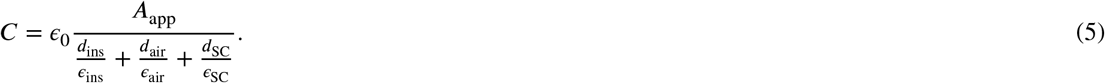

Combining Eqs. 4 and 5 gives

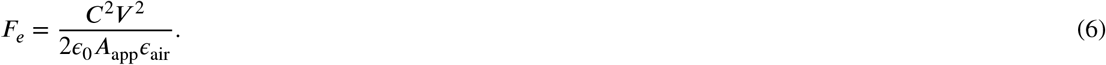

Eq. 6 includes *V*, which accounts for the voltage drop across the known environmental impedance introduced by the grounding wristband and the shunt resistor in our setup. Because of this series contribution, the nominal input voltage does not act entirely across the finger–touchscreen interaction. Following an impedance-correction approach similar to [33], the effective voltage amplitude acting across the interaction region was estimated using a magnitude-level voltage-division approximation:

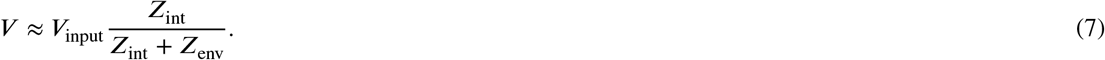

In Eq. 7, is used to denote the magnitude of the complex interaction impedance, *Z*_int_ = |*Z*_int_(*f*_0_)|, obtained after removing the known environmental contribution from the measured total impedance as in Eqs. 1 and 2. Thus, Eq. 7 provides a magnitude-level estimate of the fraction of the nominal voltage acting across the finger–touchscreen interaction, following [33]. When *Z*_env_ ≪ *Z*_int_, Eq. 7 reduces to *V* ≈ *V*_input_.

In addition, the effective gap thickness, referred to here as the total thickness between the dielectric layers, was derived from the measured effective capacitance and apparent contact area as *d*_eff_ = *ε*_0_*ε*_eff_ *A*_app_/*C*_eff_. Following prior work, which suggested that during sliding the air-gap impedance can dominate the overall electrical impedance [14, 23], we used an air-like effective permittivity, *ε*_eff_ = *ε*_air_ [23]. The resulting *d*_eff_ should not be interpreted as a direct physical measurement of the air-gap thickness. Instead, it represents an air-equivalent effective separation inferred from *C*_eff_ and *A*_app_, combining the effective contributions of the insulating layer, fingertip–surface separations, and the stratum corneum into a single air-equivalent distance.

#### Physics-informed electrostatic attraction force model

We formulated a physics-informed electrostatic attraction model to describe how normal force and sliding speed affect the electrostatic attraction from contact area and electrical impedance (Fig. 2). The model is based on the parallel-plate relation in Eq. 6. Specifically, the capacitance term was represented by the effective capacitance, *C*_eff_, obtained from the interaction admittance, effective voltage, *V*, and the apparent contact area, *A*_app_, from FTIR fingerprint images.

Although normal force and sliding speed do not appear explicitly in Eq. 6, their effects can be incorporated through the measured contact area and electrical impedance. Therefore, we treated *C*_eff_, *A*_app_, and *V* as condition-dependent quantities, such that the force–speed dependence of electrostatic attraction is represented through their dependence on normal force and sliding speed. This gives

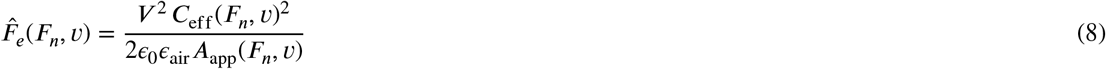

where *V* is approximated using the magnitude of the interaction impedance as 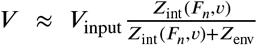. Here, *V*_input_ = *V*_rms_ was used because the experimentally derived electrostatic attraction represents a cycle-averaged force. This approach allowed the model to include the systematic effects of normal force and sliding speed on the electrostatic attraction. The specific parameterization of these condition-dependent quantities and the corresponding model evaluation are presented in the Results section.

## 3. Results

Under actuation, both the measured real contact area and the tangential force oscillated at twice the drive frequency, consistent with the quadratic dependence of electrostatic attraction on voltage [15, 24, 27, 34, 35] (Supplementary Text, Fig. S7). Consequently, the measured cycle-averaged real contact area and tangential force were higher when voltage was on compared to when it was off (Fig. 3A and Figs. 3C–E), in agreement with previous studies [10, 22, 24, 27, 35, 36, 37]. Consistently, fingerprint images showed darker regions—indicating larger optically resolved real contact area—when voltage was applied (Figs. 3A–B, Movies S1 and S2).

**Figure 3:**
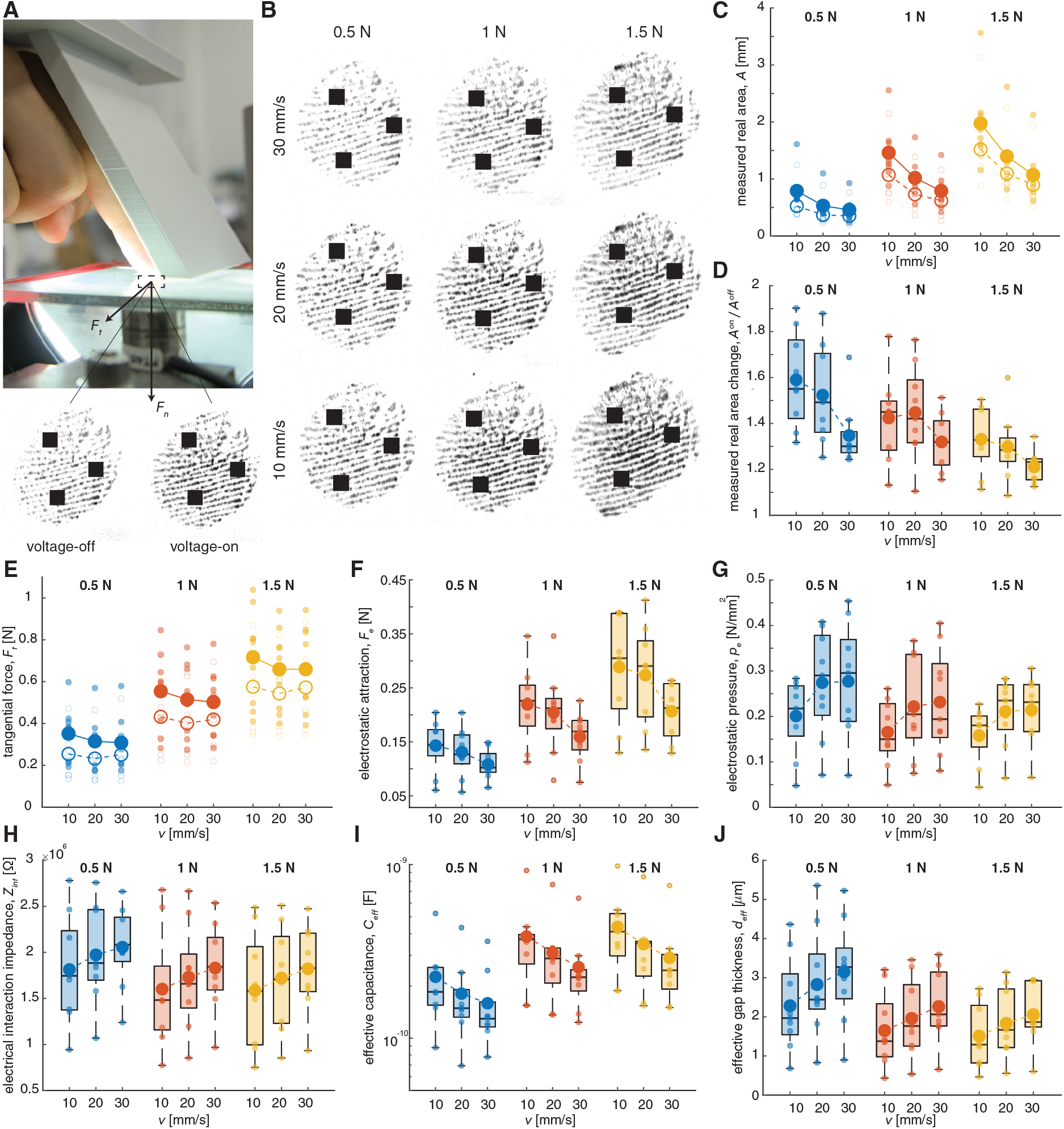
Experimental results. (A) FTIR-based imaging setup for capturing contact area during sliding; example images are shown for voltage-off and voltage-on conditions. (B) Representative FTIR-derived real-contact images across the nine touch conditions. (C) Measured real contact area *A* and (D) real-area ratio *A*^on^/*A*^off^ during sliding. (E) Tangential force *F*_*t*_, (F) electrostatic attraction force *F*_*e*_, (G) electrostatic pressure *p*_*e*_, and (H) electrical interaction impedance *Z*_int_, (I) effective capacitance *C*_eff_, and (J) effective gap thickness *d*_eff_ as functions of sliding speed and normal force. Small circles denote individual participants (averaged over repetitions within each condition); large circles indicate across-participant means (filled circles: voltage-on; open circles: voltage-off).

Overall, visual inspection of the data reveals distinct trends with normal force and sliding speed (Figs. 3C–3J). As the normal force increases, the measured real contact area, tangential force, electrostatic attraction, and effective capacitance increase, whereas *A*^on^/*A*^off^, electrostatic pressure, electrical interaction impedance, and effective gap thickness decrease. In contrast, as sliding speed increases, electrostatic pressure, electrical interaction impedance, and effective gap thickness increase, while the measured real contact area, tangential force, *A*^on^/*A*^off^, electrostatic attraction, and effective capacitance decrease.

To quantify the effects of normal force, sliding speed, and participant-level variability, we fitted linear mixed-effects models (LMMs) for each measured and derived outcome: measured real contact area, real-area change (*A*^on^/*A*^off^), tangential force, electrostatic attraction, electrostatic pressure, electrical interaction impedance, effective capacitance, and effective gap thickness. Speed and force were treated as continuous predictors to test linear trends, and all models used the specification *Y* ~ 1 + speed × force + (1|participant). The LMMs showed that force and speed affected the different contact, electrical, and electrostatic quantities in distinct ways, while participant-level variability was significant in all models.

The LMM results confirmed distinct force–speed dependencies across the measured and derived outcomes (Table 1). Measured real contact area, tangential force, electrostatic attraction, and effective capacitance increased significantly with normal force, whereas *A*^on^/*A*^off^, electrical interaction impedance, and effective gap thickness decreased with force. Sliding speed significantly increased electrostatic pressure, electrical interaction impedance, and effective gap thickness, while reducing the actuation-induced area change. Significant speed-by-force interactions were observed for measured real contact area and electrostatic attraction, indicating that their force dependence weakened at higher speeds. In all models, including a participant-specific random intercept significantly improved model fit compared with the fixed-effects-only model (likelihood-ratio tests, all *p* < 0.001).

**Table 1.**
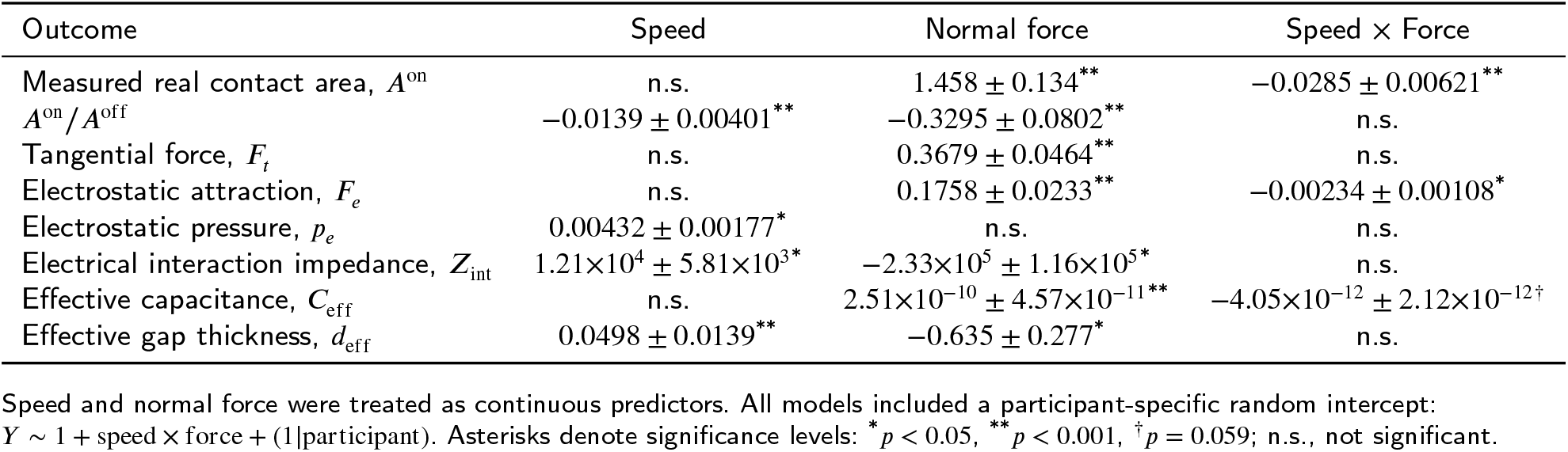
Linear mixed-effects model results for the effects of sliding speed, normal force, and their interaction. Values indicate fixed-effect estimates *β* ± *SE*.

| Outcome | Speed | Normal force | Speed × Force |
| --- | --- | --- | --- |
| Measured real contact area, $A^{\text{on}}$ | n.s. | $1.458 \pm 0.134^{**}$ | $-0.0285 \pm 0.00621^{**}$ |
| $A^{\text{on}}/A^{\text{off}}$ | $-0.0139 \pm 0.00401^{**}$ | $-0.3295 \pm 0.0802^{**}$ | n.s. |
| Tangential force, $F_t$ | n.s. | $0.3679 \pm 0.0464^{**}$ | n.s. |
| Electrostatic attraction, $F_e$ | n.s. | $0.1758 \pm 0.0233^{**}$ | $-0.00234 \pm 0.00108^*$ |
| Electrostatic pressure, $p_e$ | $0.00432 \pm 0.00177^*$ | n.s. | n.s. |
| Electrical interaction impedance, $Z_{\text{int}}$ | $1.21 \times 10^4 \pm 5.81 \times 10^3^*$ | $-2.33 \times 10^5 \pm 1.16 \times 10^5^*$ | n.s. |
| Effective capacitance, $C_{\text{eff}}$ | n.s. | $2.51 \times 10^{-10} \pm 4.57 \times 10^{-11}^{**}$ | $-4.05 \times 10^{-12} \pm 2.12 \times 10^{-12}^\dagger$ |
| Effective gap thickness, $d_{\text{eff}}$ | $0.0498 \pm 0.0139^{**}$ | $-0.635 \pm 0.277^*$ | n.s. |

Speed and normal force were treated as continuous predictors. All models included a participant-specific random intercept: $Y \sim 1 + \text{speed} \times \text{force} + (1|\text{participant})$ . Asterisks denote significance levels: $^*p < 0.05$ , $^{**}p < 0.001$ , $^\dagger p = 0.059$ ; n.s., not significant.

We further examined correlations among the measured outcomes by quantifying pairwise associations using three complementary Spearman correlations: (i) *pooled*, computed across all observations; (ii) *partial*, obtained after regressing out sliding speed and normal force (i.e., association at matched speed and force); and (iii) *within-participant*, computed after demeaning each variable within participant to quantify co-variation across conditions within individuals. The resulting correlation patterns were separated into robust associations (measured real contact area vs. tangential force; electrical interaction impedance vs. electrostatic pressure) and summary-dependent associations involving electrostatic attraction.

We observed a strong positive association between measured real contact area and tangential force, consistent with previous studies [4, 24] (Fig. 4A; pooled *r* = 0.75, *p* < 0.001; partial *r* = 0.59, *p* < 0.001; within-participant *r* = 0.86, *p* < 0.001). In contrast, the association between electrostatic attraction and measured real contact area depended on how shared variation due to force and speed was handled (pooled *r* = 0.48, *p* < 0.001; partial *r* = −0.16, *p* = 0.166; within-participant *r* = 0.86, *p* < 0.001). Similarly, the association between *A*^on^/*A*^off^ and electrostatic attraction varied across summaries (Fig. 4B; pooled *r* = 0.28, *p* < 0.05; partial *r* = 0.74, *p* < 0.001; within-participant *r* = −0.2, *p* = 0.07). We examined the association between *A*^on^/*A*^off^ and electrostatic attraction under different force conditions. When the normal force was fixed, larger actuation-induced area increases were consistently associated with larger inferred electrostatic attraction. The pooled correlation coefficients were *r* = 0.69 (*p* < 0.001) for 0.5 N, pooled *r* = 0.74 (*p* < 0.001) for 1 N, and pooled *r* = 0.95 (*p* < 0.001) for 1.5 N. Finally, electrical interaction impedance and electrostatic pressure were consistently positively correlated (Fig. 4C; pooled *r* = 0.5, *p* < 0.001; partial *r* = 0.46, *p* < 0.001; within-participant *r* = 0.63, *p* < 0.001).

**Figure 4:**
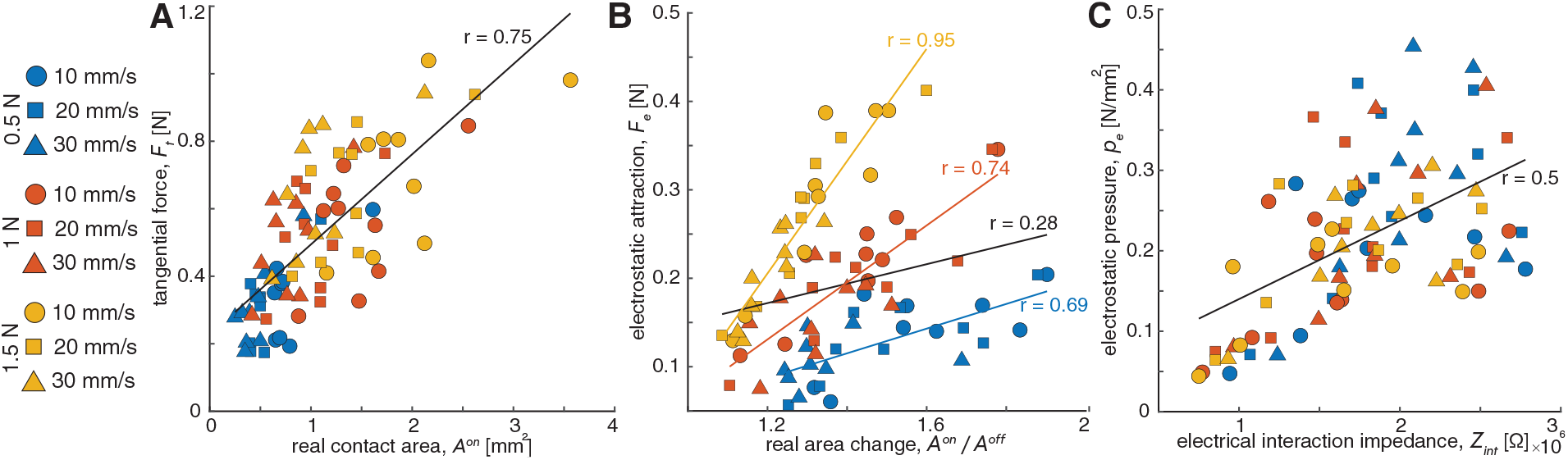
Correlations between measured outcomes. (A) Tangential force versus real contact area, showing a strong positive association. (B) Electrostatic attraction versus real-area ratio *A*^on^/*A*^off^, indicating that larger voltage-induced increases in contact area correspond to higher attraction. (C) Electrostatic pressure versus electrical interaction impedance, revealing a positive association across conditions. Colors indicate normal force (0.5, 1.0, and 1.5 N) and marker shape indicates sliding speed (10, 20, and 30 mm/s). *r* denotes the pooled Spearman correlation.

### 3.1. Physics-informed modeling of electrostatic attraction

After establishing systematic force–speed effects and correlations among the measured contact area and electrical impedance quantities, we next evaluated whether these quantities could be combined within the physics-informed framework to estimate electrostatic attraction. Because electrostatic attraction varied systematically with normal force and sliding speed, we used the parallel-plate-based formulation presented in Section 2.5. The results are summarized in Fig. 2, showing how touch conditions map to electrostatic attraction via measurable contact quantities.

#### Effective capacitance, apparent contact area, and electrical interaction impedance models

The intermediate quantities used in the electrostatic attraction model, namely the effective capacitance, *C*_eff_, apparent contact area, *A*_app_, and magnitude of the electrical interaction impedance, *Z*_int_, varied across the nine force–speed conditions. To use these measured quantities as inputs to the electrostatic attraction model, we first expressed their variation across the tested force–speed conditions as compact functions of normal force, *F*_*n*_, and sliding speed, *v*.

For each force–speed condition, mean-level values were obtained by averaging the corresponding measured quantity across participants. These condition-averaged values were then fitted using power-law functions of normal force and sliding speed. For example, the effective capacitance was represented as

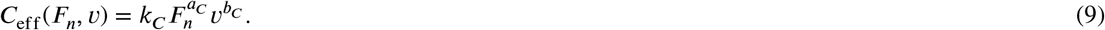

The coefficients were obtained by linear regression in the log-transformed domain,

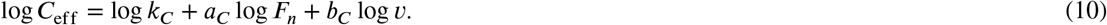

The same log–log fitting procedure was used to describe the force–speed dependence of the apparent contact area,

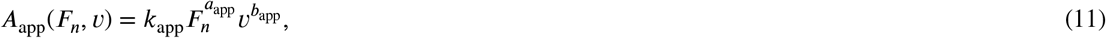

and the magnitude of the interaction impedance,

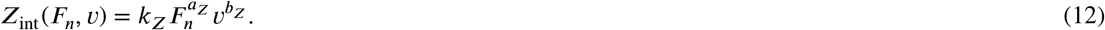

Details of the fitting procedure are provided in the Supplementary Text.

The fitted power-law parameters and error metrics are summarized in Table 2. The low normalized errors indicate that these mean-level functions captured the main force–speed dependence of the intermediate contact area and electrical impedance quantities well (Figs. 5A–C), and were therefore used as inputs to the electrostatic attraction model. Additional mean-level parameterizations for the measured real contact area and the effective gap thickness are provided in the Supplementary Information (Fig. S8).

**Table 2.** Mean-level power-law model parameters and fit errors. Each quantity was fitted as 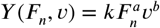 with *F*_*n*_ in N and *v* in m/s.

| $Y$ | $k$ | $a$ | $b$ | $R^2$ | RMSE | MAE | NRMSE |
| --- | --- | --- | --- | --- | --- | --- | --- |
| $C_{eff}$ [F] | $7.12 \times 10^{-11}$ | 0.601 | -0.351 | 0.96 | $1.79 \times 10^{-11}$ | $1.53 \times 10^{-11}$ | 0.064 |
| $A_{app}$ [m<sup>2</sup>] | $5.05 \times 10^{-5}$ | 0.180 | -0.015 | 0.97 | $8.00 \times 10^{-7}$ | $6.47 \times 10^{-7}$ | 0.070 |
| $Z_{int}$ [Ω] | $2.86 \times 10^6$ | -0.125 | 0.120 | 0.94 | $3.45 \times 10^4$ | $3.19 \times 10^4$ | 0.074 |

**Figure 5:**
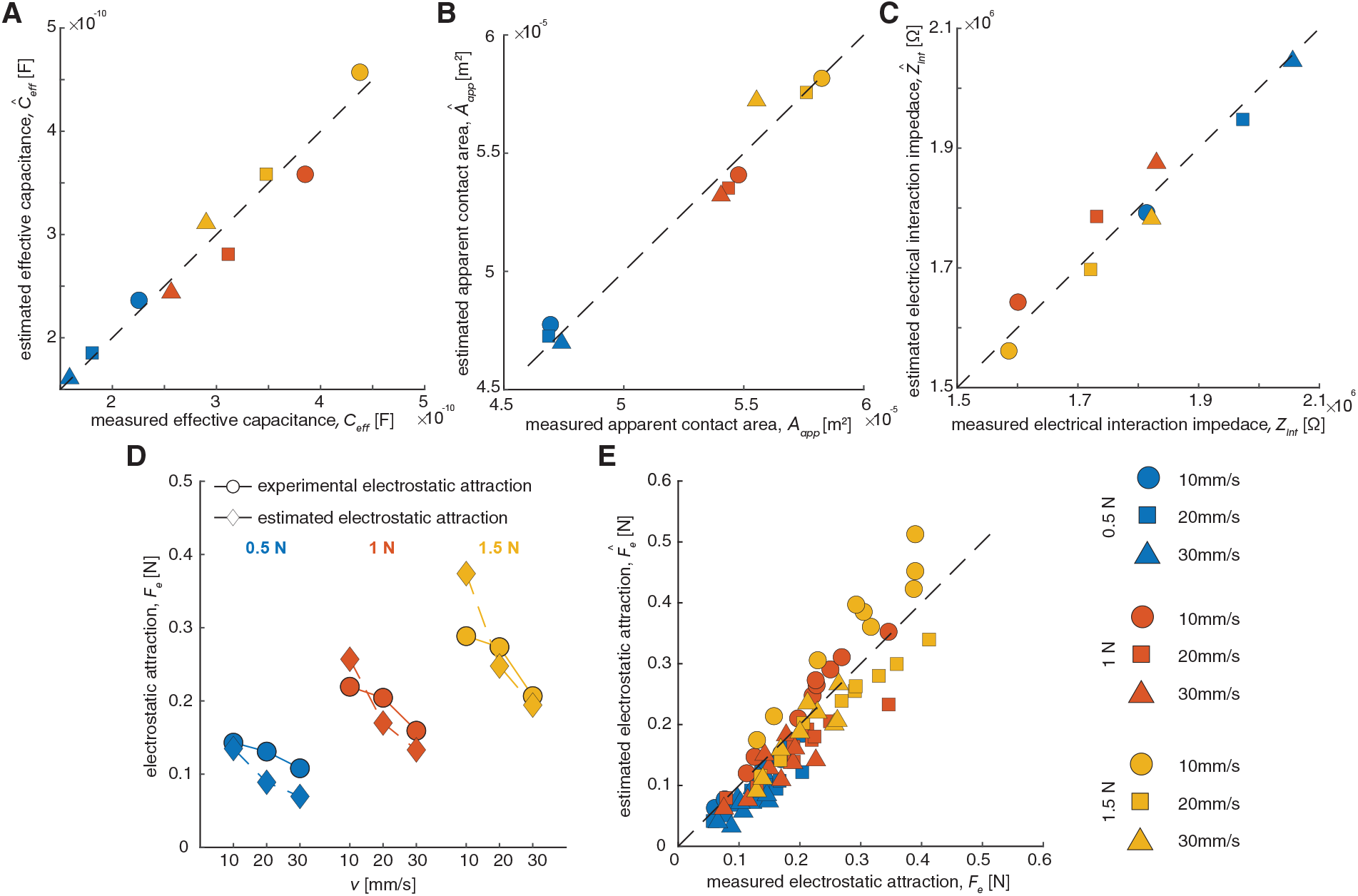
Modeling results. (A–C) Measured-versus-estimated comparisons for the mean-level power-law parameterizations of effective capacitance *C*_eff_, apparent contact area *A*_app_, and electrical interaction impedance *Z*_int_. (D) Condition-wise comparison between measured and estimated mean-level electrostatic attraction. Circles indicate measured condition means and diamonds indicate model estimates. (E) Participant-specific model with one scaling factor per participant. Dashed lines in (A–C, E) indicate the unity line.

#### Mean-level electrostatic attraction model

The fitted mean-level functions *C*_eff_ (*F*_*n*_, *v*), *Z*_int_ (*F*_*n*_, *v*), and *A*_app_(*F*_*n*_, *v*) were inserted directly into Eq. 8 to estimate the condition-averaged electrostatic attraction force across participants. This mean-level implementation was used to evaluate whether the measured effective capacitance, interaction impedance, and apparent interaction area reproduced the correct magnitude and force- and speed-dependence of the electrostatic attraction. This yielded

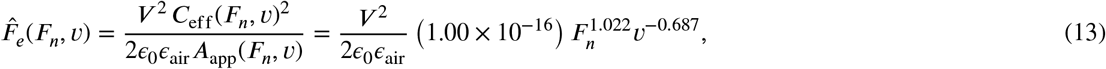

where

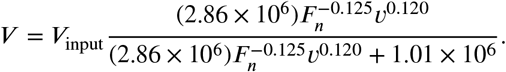

We first evaluated the mean-level implementation by using Eq. 13. This model reproduced the condition-averaged electrostatic attraction with RMSE = 0.040 N, MAE = 0.035 N, and NRMSE= 0.224 (Fig. 5D). The condition-wise comparison showed that the model captured the overall force scale and the main force–speed dependence, although deviations remained for specific force–speed conditions, particularly at 10 mm/s. These systematic deviations indicate that the idealized parallel-plate formulation does not fully account for fingertip–surface non-idealities, including possible low-speed stick–slip effects.

#### Participant-specific scaling

To account for participant-to-participant variability, we next introduced a participant-specific scaling factor to Eq. 13, while keeping the force–speed dependence fixed by the mean-level functions of *C*_eff_, *A*_app_, and *Z*_int_. The same mean-level functions were used for all participants, and a participant-specific single scaling factor *p*_1,*i*_ was determined for each participant using that participant’s measured electrostatic attraction values across the nine force–speed conditions. This factor accounts for deviations from the mean-level effective model, including participant-dependent differences in skin (electro)mechanical properties, hydration state, and contact morphology, as well as deviations from the idealized parallel-plate theory. Importantly, *p*_1,*i*_ does not introduce additional force- or speed-dependent behavior; this behavior remains fixed by the mean-level functions of *C*_eff_, *Z*_int_, and *A*_app_. The participant-specific estimated electrostatic attraction force was therefore

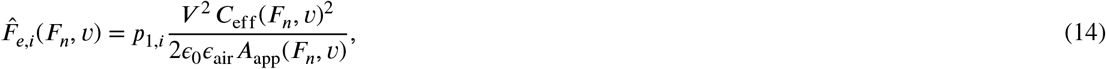

with *V* defined as in Eqs. 7 and 13.

The participant-specific scaling factor varied across participants (*p*_1,*i*_ *∈* [0.46, 1.37]), reflecting inter-participant differences in finger–screen coupling. The participant-specific model yielded RMSE = 0.045 N, MAE = 0.037 N, and NRMSE= 0.128 across the pooled participant-level data (Fig. 5E). This reduction in normalized error indicates that a single participant-specific factor accounted for part of the variability in the effective fingertip–surface contact, without modifying the force–speed dependence imposed by the physics-informed model structure.

## 4. Discussion

This study investigated how fingertip–screen contact area, interaction forces, and electrical impedance vary during fingertip sliding under different normal forces and sliding speeds. Simultaneous measurements revealed that these interaction conditions influence electrostatic attraction by modifying both the contact area and the electrical impedance. Building on these observations, we developed a physics-informed model that estimates electrostatic attraction by incorporating the measured force- and speed-dependent contact quantities. Participant-specific scaling factors further accounted for inter-participant variability arising from differences in skin (electro)mechanical properties, hydration state, contact morphology, and other non-idealities not captured by the idealized parallel-plate capacitor model [5, 21, 22, 24, 26].

The observed force- and speed-dependent changes in fingertip contact are consistent with the known mechanical behavior of the fingertip during sliding. Increasing normal force enlarges the real contact area by increasing the number and size of micro-junctions [4, 5]. Sliding speed also influences contact because of the viscoelastic nature of the fingertip. At lower speeds, the tissue has more time to relax and conform to the surface under a given load. At higher speeds, however, the reduced time available for deformation causes the tissue to behave more stiffly, leading to a smaller real contact area and a larger effective gap between the finger and the screen [13, 38, 39, 40, 41]. These observations agree with previous studies showing that apparent contact area during sliding can be smaller than under static loading at comparable normal forces [36, 38, 41, 42]. Together, these mechanical effects provide a physical basis for representing the finger–screen contact geometry as dependent on the interaction conditions in the proposed model.

The electrical response of the fingertip–screen contact also varied with the interaction conditions. Higher normal forces were associated with lower electrical interaction impedance and higher effective capacitance, whereas higher sliding speeds showed the opposite trend, with corresponding changes observed in the electrostatic attraction. The lower effective capacitance, *C*_eff_, observed at higher sliding speeds and lower normal forces may result from a smaller contact area and a larger effective separation between the fingertip and the screen [4, 5, 36, 42, 13, 41, 14, 15, 32]. Conversely, lower sliding speeds and higher normal forces may promote more sustained contact and greater moisture retention, increasing interfacial conductance and thereby reducing the electrical interaction impedance [15, 26, 43]. Supporting this interpretation, FTIR images from one participant revealed visible condensation beneath the fingerprint (Fig. S9 and Movie S3), indicating locally elevated moisture [24, 44, 45]. The same participant also exhibited lower *R*_eff_ values (Fig. S10), consistent with a higher interfacial conductance expected for a more hydrated contact [24, 26].

The equivalent air-gap thickness, estimated by combining the measurements of effective capacitance and real contact area, increased with sliding speed and decreased with normal force (Fig. 3J), consistent with the broader pattern that sliding increases air-gap thickness reported previously [14, 32]. Since the touchscreen layers and stratum corneum thickness do not change across these conditions, this trend mainly reflects changes in the air gap at the fingertip–touchscreen interface [23]. It is also consistent with fingertip conformity under load: higher normal force was associated with larger real contact area and smaller effective gap thickness [16, 24, 46, 31].

The mean-level model, which used these measured changes in apparent contact area and effective capacitance, captured the systematic force–speed trends in electrostatic attraction, supporting the interpretation that normal force and sliding speed influence electrostatic attraction through these condition-dependent contact quantities. Remaining prediction errors may arise from moisture accumulation [43], stick–slip-like fluctuations in contact area, particularly at lower sliding speeds [24], and the effects of viscoelastic finger mechanics [13, 24], which are not explicitly represented in the idealized parallel-plate model.

Nonetheless, a substantial inter-participant variability was observed in electrostatic attraction. Similar variability has been reported previously [13, 15, 24, 26] and is consistent with studies demonstrating the sensitivity of electrostatic actuation to interfacial conditions, including humidity, perspiration, contact morphology, and skin (electro)mechanical properties [8, 47, 48]. In the proposed model, this variability was represented by a participant-specific scaling factor, *p*_1,*i*_. The wide range of estimated values, *p*_1,*i*_ *∈* [0.46, 1.37], indicates that identical force–speed conditions can produce substantially different electrostatic attraction across participants. Importantly, *p*_1,*i*_ does not introduce participant-specific force or speed dependencies. Instead, the condition dependence is determined by the mean-level model through the condition-dependent apparent contact area, effective capacitance, and effective voltage, while *p*_1,*i*_ uniformly scales the predicted attraction for each participant. This approach provides a simple means of accounting for inter-participant variability while preserving a common model structure. Because *p*_1,*i*_ is a single scalar, it may also offer a practical route toward low-burden per-user calibration, although the number of trials required for reliable estimation should be investigated in future work.

The correlation analyses further highlight the importance of accounting for participant-specific differences. The relationship between measured real contact area and electrostatic attraction was strong within individual participants but weaker when all observations were pooled, indicating that inter-participant variability can obscure the condition-dependent trends observed within a given finger. One possible explanation is variation in skin hydration: moisture can reduce electrostatic attraction [24, 26] while simultaneously increasing baseline contact area and friction [24, 26, 43, 45, 47]. Consequently, across participants, a larger real contact area does not necessarily correspond to greater electrostatic attraction, even though the two quantities may vary together within the same participant. Accordingly, normalized measures such as the real-area ratio, *A*^on^/*A*^off^, can help separate actuation-induced contact changes from baseline differences between participants [24, 36].

Despite careful experimental design and analysis, this study has several limitations. First, the FTIR-based imaging and analysis method [29] provides an optically resolved measure of fingertip–glass contact area and may not capture the absolute real contact area at microscopic or nanoscopic length scales. As real contact area is inherently scale-dependent, unresolved surface roughness can influence its absolute magnitude [49]. Consequently, the contact area reported here should be interpreted as an optically resolved contact metric rather than the full multiscale real contact area. Second, the proposed model may be less applicable under excessive moisture conditions, where increased interfacial conductance and other non-idealities can violate the assumptions of the parallel-plate representation. Third, the effective capacitance and equivalent air-gap thickness values were estimated from impedance measurements at a single actuation frequency (75 Hz) and should therefore be interpreted as frequency-specific effective parameters. Although this frequency was selected to minimize the influence of the experimental setup and lies within a perceptually relevant range for electrostatic stimulation, both the electrical behavior of the fingertip–screen interface (e.g., charge relaxation and leakage) and the mechanical response of the fingertip are frequency dependent [13, 18, 24, 32, 31]. Accordingly, the proposed model should be interpreted as describing the effects of normal force and sliding speed at the tested stimulation frequency rather than as a frequency-general model of electrostatic attraction. Finally, we did not directly measure participant-specific skin properties, such as hydration, stiffness, curvature, or fingerprint morphology, which may contribute to the observed inter-participant variability. Although these factors are implicitly captured by the lumped scaling factor *p*_1,*i*_, this parameter does not distinguish their individual physiological, mechanical, or morphological contributions. Future work combining optically resolved contact imaging with direct measurements of fingertip properties and broader frequency-dependent measurements could further clarify the physical sources of inter-participant variability and extend the model to a broader range of stimulation frequencies.

## 5. Conclusion

In summary, this study shows that electrostatic attraction during fingertip sliding depends on how the fingertip– screen contact changes with normal force and sliding speed. The simultaneous measurements of optically resolved contact area, interaction forces, and electrical impedance revealed that contact area, tangential force, electrical interaction impedance, effective capacitance, effective voltage, and effective gap thickness varied jointly and systematically across the tested force–speed conditions. By incorporating these force- and speed-dependent variables, a physics-informed parallel-plate capacitor model was proposed to estimate electrostatic attraction. The model reproduced the condition-averaged electrostatic attraction, while participant-specific scaling factors accounted for inter-participant variability.

Together, these findings provide an experimentally grounded framework for estimating electrostatic attraction under realistic touch conditions. The results support the design of condition- and user-aware electrostatic haptic interfaces, where voltage modulation can be adapted not only to the desired tactile effect but also to the user’s normal force, sliding speed, and contact state. Moreover, the proposed model provides a foundation for adaptive control strategies that use fingertip–surface contact area and electrical impedance measurements to regulate electrostatic feedback across users and interaction conditions.

## 8. Data availability

The data used in this paper will be publicly available upon acceptance.

## 9. AI usage

The authors used OpenAI’s ChatGPT and Grammarly Inc.’s Grammarly for language editing and readability improvements. They are responsible for all content and the final manuscript.

### CRediT authorship contribution statement

#### Celal Umut Kenanoglu

Conceptualization, Methodology, Investigation, Software, Hardware, Formal Analysis, Data Curation, Visualization, Writing - original draft. **Yasemin Vardar:** Conceptualization, Methodology, Formal Analysis, Visualization, Writing - original draft, Writing - review & editing, Supervision, Resources, Project Administration, Funding Acquisition.

