## Supplementary Information for "Physics-Informed Estimation of Electrostatic Attraction During Fingertip Sliding Under Varying Speed and Normal Force"

### Supplementary Text

#### Materials and Methods

##### Data Acquisition and Extraction

Fig. S1 illustrates the experimental setup; full details are provided in the Materials and Methods section of the main text.

Fig. S4a depicts the tangential force signals recorded during sliding in both directions (left-to-right and right-to-left). As the sign of tangential force changes with sliding direction, we also report the absolute tangential force computed only within the camera-trigger window; these values are pooled across all trials that passed the validity criteria and are summarized in Fig. S4b. Fig. S4c provides a zoomed segment from a representative trial, highlighting the periodic (sinusoidal) modulation of tangential force observed under the applied voltage. Importantly, the local minima of this modulation occur at the zero-crossings of the voltage signal ($V = 0$ V) and coincide with the tangential force levels measured in voltage-off segments, indicating that the two ways of sampling the "no-voltage" state are consistent.

Camera acquisition was initiated by a trigger sent from MATLAB/Simulink to the Motion-BLITZ software, after which frames were recorded at 1000 frames per second (fps). The voltage-on timing marker in Fig. S4a was used to define the analysis window and to segment voltage-on and voltage-off intervals from within the same trial, thereby aligning force and image-based quantities in time. The raw FTIR images could not be analyzed directly due to geometric warping caused by (i) lens-related radial distortion and (ii) the oblique camera viewing angle relative to the glass surface. We therefore applied a two-step correction. First, radial distortion was compensated using intrinsic parameters obtained from a checkerboard calibration. Second, we applied a projective transformation [1] to obtain a virtual top-down view of the contact region. The transformation matrix was determined using an image of a circular rubber reference placed on the glass: the observed ellipse was mapped to an ideal circle. Real contact area was then computed from the corrected FTIR images following [2].

The resulting real contact area during the camera-trigger interval is reported in Fig. S5a. A representative trace from 20 mm/s is shown in Fig. S5b, where contact area varies periodically in time. As with tangential force, the troughs in the real contact area trace align with the voltage zero-crossings ($V = 0$ V) and match the values measured during voltage-off segments, supporting consistency between the two states. As discussed in the main text, the increase in real contact area is attributed to the pulling effect of the electrostatic attraction.

Fig. S6 shows a representative current trace from a single trial. After removing the DC component, the signal appears as a near-sinusoid, with its dominant oscillation occurring at the electrostatic actuation frequency.

#### Parallel-plate Capacitive Theory - Voltage doubling

The normal electrostatic force (electrostatic attraction) at the fingertip–screen interface can be calculated as follows [3, 4, 5] as presented in main text:

$$F_e = \epsilon_0 \frac{A_{\text{app}} V^2}{2\left(\frac{d_{\text{ins}}}{\epsilon_{\text{ins}}} + \frac{d_{\text{air}}}{\epsilon_{\text{air}}} + \frac{d_{\text{SC}}}{\epsilon_{\text{SC}}}\right)^2 \epsilon_{\text{air}}}, \tag{1}$$

Because the electrostatic force is proportional to the square of the applied voltage, applying a zero-mean sinusoid $V = V_0 \sin(\omega t)$ gives

$$V^2 = V_0^2 \frac{1 - \cos(2\omega t)}{2}$$

using $\sin^2(\omega t) = \frac{1-\cos(2\omega t)}{2}$. This form makes the $2\omega$ term explicit and therefore predicts a frequency-doubling component in the electrostatic force. Consistent with this nonlinear voltage–force dependence in Eq. 1, the Fourier spectrum of the real contact area (Fig. S7b) shows a dominant peak at twice the electrostatic actuation frequency, similar to the peak observed for the tangential force in Fig. S7a. The Fourier spectrum of the current signal (Fig. S7c) shows a clear peak at the electrostatic actuation frequency, as expected for a measurement that directly follows the voltage drive applied to the touchscreen.

### Power-law Parameterization

To describe the dependence of the measured electrical and contact quantities on normal force and sliding speed, we used empirical power-law models of the form

$$Y(F_n, v) = k_Y F_n^{a_Y} v^{b_Y}, \tag{2}$$

where $Y$ denotes the quantity of interest, $F_n$ is the normal force [N], and $v$ is the sliding speed [m/s]. For each variable, the model parameters were obtained by performing a linear regression in the log–log domain:

$$\log Y = \beta_0 + \beta_1 \log F_n + \beta_2 \log v. \tag{3}$$

The coefficients in the original domain were then calculated as $k_Y = e^{\beta_0}$, $a_Y = \beta_1$, and $b_Y = \beta_2$. The goodness of fit was evaluated over the nine tested $(F_n, v)$ conditions using the coefficient of determination, RMSE, MAE, and NRMSE. Unless otherwise stated, NRMSE was computed by normalizing the RMSE by the range of the measured values.

#### Measured real contact area model $A^{\text{on}}(F_n, v)$

The same procedure was applied to the mean measured real contact area under electrostatic actuation. Let $A^{\text{on}}$ [m$^2$] denote the measured real contact area for each condition. The log–log model was

$$\log A^{\text{on}} = \beta_0 + \beta_1 \log F_n + \beta_2 \log v. \tag{4}$$

The fitted power-law model was

$$A^{\text{on}}(F_n, v) = k_A F_n^{a_A} v^{b_A}, \quad k_A = 1.21 \times 10^{-7},\ a_A = 0.834,\ b_A = -0.537. \tag{5}$$

The model yielded $R^2 = 0.997$, RMSE$= 2.48 \times 10^{-8}$ m$^2$, MAE$= 2.19 \times 10^{-8}$ m$^2$, and NRMSE$=$ 0.016 over the nine conditions. These coefficients indicate that the real contact area increases strongly with normal force and decreases with sliding speed (Fig. S8a).

#### Effective gap thickness model $d_{\text{eff}}(F_n, v)$

The effective gap thickness was treated as a derived quantity rather than an independently measured variable. It was obtained from the measured electrical and contact quantities and then modeled as a function of normal force and sliding speed. Let $d_{\text{eff}}$ [m] denote the effective gap thickness for each condition. The log–log regression was

$$\log d_{\text{eff}} = \beta_0 + \beta_1 \log F_n + \beta_2 \log v. \tag{6}$$

The fitted model was

$$d_{\text{eff}}(F_n, v) = k_d F_n^{a_d} v^{b_d}, \quad k_d = 6.43 \times 10^{-6},\ a_d = -0.398,\ b_d = 0.287. \tag{7}$$

The model yielded $R^2 = 0.980$, RMSE$= 7.14 \times 10^{-8}$ m, MAE$= 6.53 \times 10^{-8}$ m, and NRMSE$=$ 0.006 over the nine conditions. The negative force exponent indicates that the effective gap decreases with increasing normal force, while the positive speed exponent indicates that the effective gap increases with sliding speed. This trend is consistent with the observed decrease in real contact area and capacitance at higher sliding speeds (Fig. S8b).

#### Capacitance model $C_{\text{eff}}(F_n, v)$

We modeled the mean effective capacitance across participants as a power law of normal force and sliding speed. Let $C_{\text{eff}}$ [F] denote the measured mean capacitance for each of the nine $(F_n, v)$ conditions. The log–log regression was written as

$$\log C_{\text{eff}} = \beta_0 + \beta_1 \log F_n + \beta_2 \log v. \tag{8}$$

The resulting model in the original domain was

$$C_{\text{eff}}(F_n, v) = k_C F_n^{a_C} v^{b_C}, \quad k_C = 7.12 \times 10^{-11},\ a_C = 0.601,\ b_C = -0.351. \tag{9}$$

The model yielded $R^2 = 0.958$, RMSE$= 1.79 \times 10^{-11}$ F, MAE$= 1.53 \times 10^{-11}$ F, and NRMSE$=$ 0.064 over the nine conditions. The positive force exponent and negative speed exponent indicate that the effective capacitance increases with normal force and decreases with sliding speed over the tested range.

#### Apparent contact area model $A_{\text{app}}(F_n, v)$

The same log–log fitting procedure was applied to the mean apparent contact area. Let $A_{\text{app}}$ [m$^2$] denote the apparent contact area for each of the nine conditions. The regression model was

$$\log A_{\text{app}} = \beta_0 + \beta_1 \log F_n + \beta_2 \log v. \tag{10}$$

The resulting model was

$$A_{\text{app}}(F_n, v) = k_{\text{app}} F_n^{a_{\text{app}}} v^{b_{\text{app}}}, \quad k_{\text{app}} = 5.05 \times 10^{-5},\ a_{\text{app}} = 0.180,\ b_{\text{app}} = -0.015. \tag{11}$$

The model yielded $R^2 = 0.966$, RMSE$= 8.00 \times 10^{-7}$ m$^2$, MAE$= 6.47 \times 10^{-7}$ m$^2$, and NRMSE$=$ 0.070 over the nine conditions. The small velocity exponent, $b_{\text{app}} = -0.015$, indicates that the apparent contact area is only weakly affected by sliding speed over the tested range. Therefore, $A_{\text{app}}$ is mainly described by its dependence on normal force, while the speed dependence is negligible compared with the measured real contact area.

#### Electical interaction impedance model $Z_{\text{int}}(F_n, v)$

The same procedure was applied to the mean electrical interaction impedance. Let $Z_{\text{int}}$ [$\Omega$] denote the measured mean impedance for each of the nine $(F_n, v)$ conditions. The log–log model was

$$\log Z_{\text{int}} = \beta_0 + \beta_1 \log F_n + \beta_2 \log v. \tag{12}$$

The fitted model in the original domain was

$$Z_{\text{int}}(F_n, v) = k_Z F_n^{a_Z} v^{b_Z}, \quad k_Z = 2.86 \times 10^6,\ a_Z = -0.125,\ b_Z = 0.120. \tag{13}$$

The model yielded $R^2 = 0.944$, RMSE$= 3.45 \times 10^4\ \Omega$, MAE$= 3.19 \times 10^4\ \Omega$, and NRMSE$= 0.074$ over the nine conditions. The negative force exponent and positive velocity exponent indicate that the electrical impedance decreases with increasing normal force and increases with sliding speed. These trends are consistent with improved interfacial coupling at higher normal forces and reduced effective contact at higher sliding speeds.

#### Summary of fitted power-law models and additional results

Overall, the fitted models captured the dependence of the measured electrical and contact quantities on normal force and sliding speed with high goodness of fit across the nine experimental conditions. The real contact area and capacitance both increased with normal force and decreased with sliding speed, whereas the effective gap thickness and electrical impedance showed the opposite trend with speed. The apparent contact area showed a much weaker dependence on sliding speed, suggesting that the speed-dependent changes observed in the electrical response are more closely related to changes in real contact area and effective interfacial gap than to changes in the macroscopic apparent area.

To characterize the mechanical response of the experimental setup, an impact-hammer test was performed on the touchscreen and force-sensor assembly. The setup was excited using an instrumented impact hammer in the tangential direction, and the corresponding tangential force-sensor response was used to estimate the frequency response of the system. Because electrostatic attraction scales with the square of the applied voltage, a 75 Hz sinusoidal voltage is expected to produce a dominant force component at 150 Hz. Therefore, the setup response was inspected around both the selected voltage frequency of 75 Hz and the expected force-response frequency of 150 Hz. As shown in Fig. S2, the response around the selected stimulation frequency of 75 Hz was relatively stable and did not coincide with a pronounced resonance peak. This characterization supported the use of 75 Hz in the sliding experiments, reducing the potential influence of setup dynamics on the measured signal.

Frequency-dependent characterization was performed using the same participant group under a fixed sliding condition of $F_n = 1$ N and $v = 20$ mm/s [6]. A sinusoidal voltage of 100 $\text{V}_{\text{peak}}$ was applied while the stimulation frequency was varied logarithmically between 25 and 2500 Hz. As shown in Fig. S3a, $C_{\text{eff}}$ was high at low frequencies and decreased strongly in the low-frequency range, varied only modestly over the intermediate frequency range, including 75 Hz, and showed a slight decrease at the very high frequencies. This trend may reflect the lossy nature of the fingertip–surface contact, where slow charge redistribution, interfacial polarization, and moisture-related pathways can affect the admittance-derived capacitance at low frequencies. At higher frequencies, these slower polarization processes may no longer follow the rapidly alternating field [7], reducing their contribution to $C_{\text{eff}}$. One likely reason is that the parallel-plate capacitance representation assumes an idealized capacitive coupling, whereas the real fingertip–surface contact is lossy and can involve resistive pathways, charge relaxation, leakage, and moisture-related interfacial effects [8, 6, 9, 10]. In addition, the friction-derived electrostatic attraction may include frequency-dependent mechanical and tribological transfer effects [6, 11], which are not captured by $C_{\text{eff}}$ alone.

Fig. S9 shows a moist finger, where condensation is observed behind the fingertip. For this fingertip, the $R_{\text{eff}}$ value was the lowest. The values of $R_{\text{eff}}$ is presented in Fig. S10. $R_{\text{eff}}$ increases with higher speed and lower force.

Representative recordings of fingertip contact are presented in Movies S1–S3. Movies S1 and S2 illustrate different touch conditions, whereas Movie S3 compares dry and moist finger states. No post-processing corrections—such as optical distortion or angular adjustments—were applied to these recordings in order to preserve their anonymity.

### Supplementary Figures

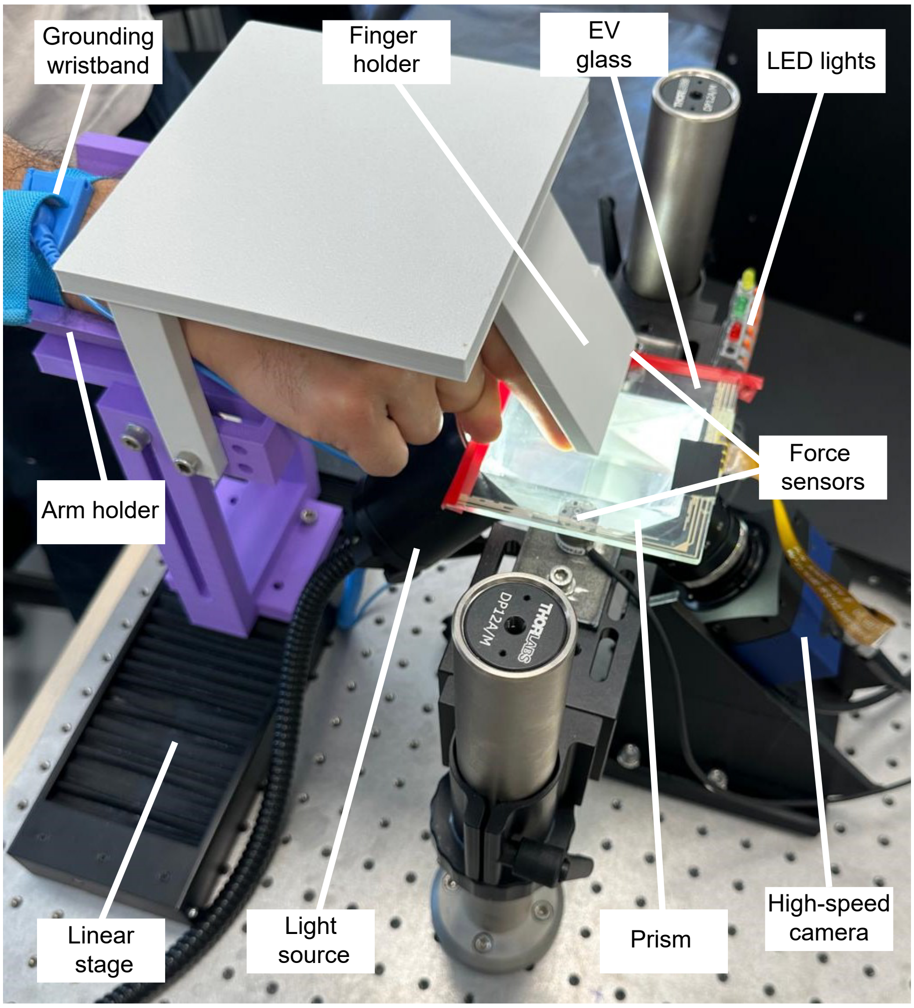


**Figure S1: Experimental setup.**

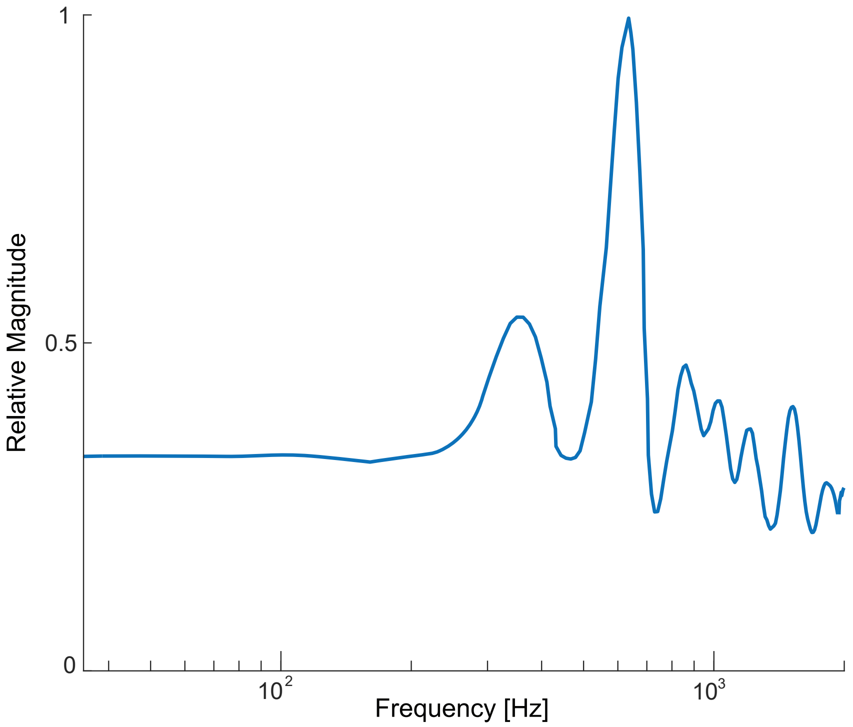


**Figure S2: Impact-hammer test result of the experimental setup in the tangential direction.**

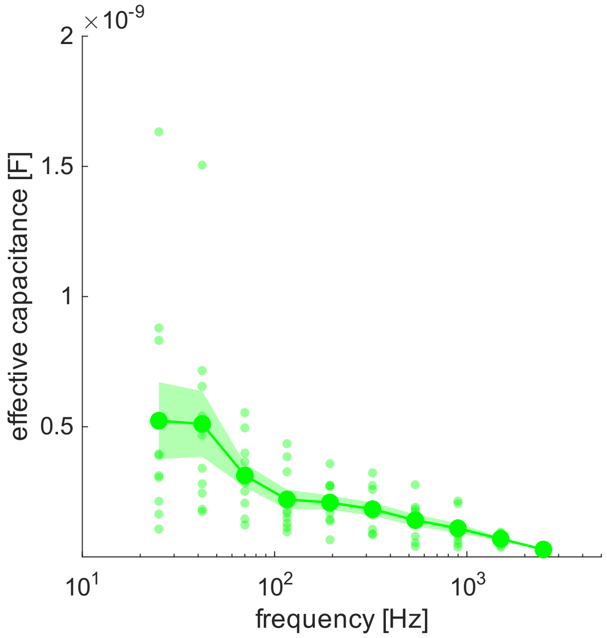


**(a)**

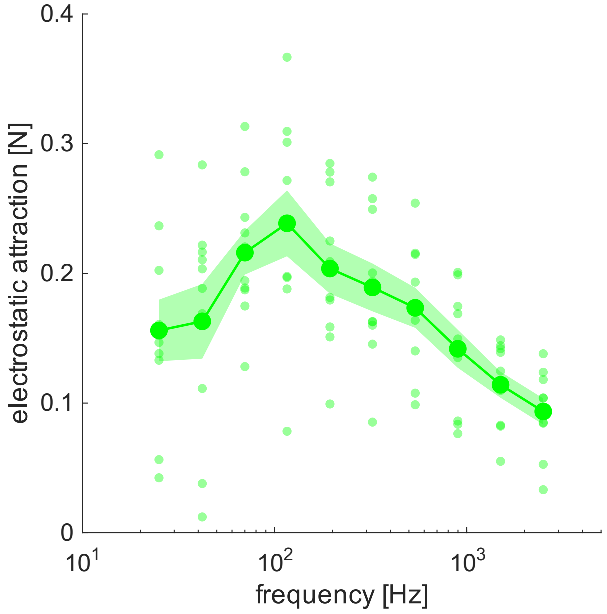


**(b)**

**Figure S3: Frequency-dependent effective capacitance and electrostatic attraction.** (a) Effective capacitance ($C_{\mathrm{eff}}$) as a function of stimulation frequency. (b) Friction-derived electrostatic attraction as a function of stimulation frequency. Light green points indicate individual measurements, solid markers indicate mean values, and shaded regions indicate variability across measurements.

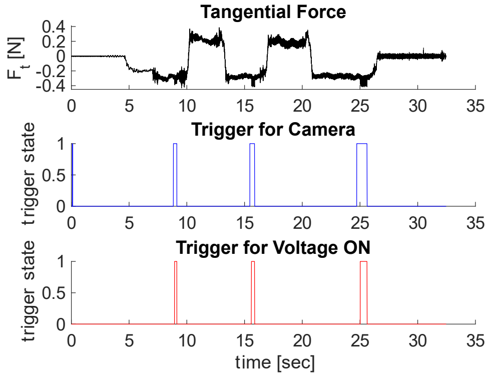


(a)

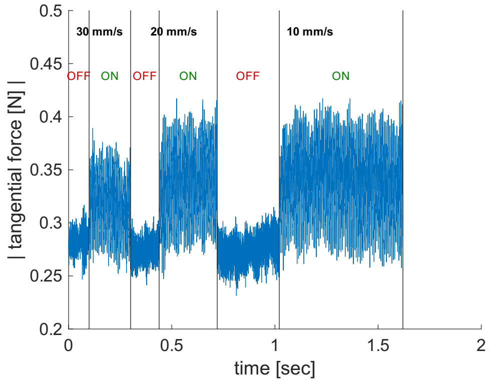


(b)

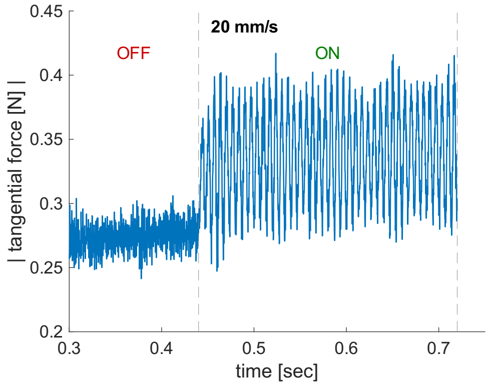


(c)

**Figure S4: Extracted tangential force and trigger signals from Participant-1 when the applied normal force was 1 N.** (a) Raw tangential force data recorded during a trial with voltage and camera trigger signals. Values of 1 and 0 indicate that the trigger is active and inactive, respectively. (b) Absolute value of tangential force data when the camera trigger is on. (c) Absolute value of tangential force measurement for 20 mm/s.

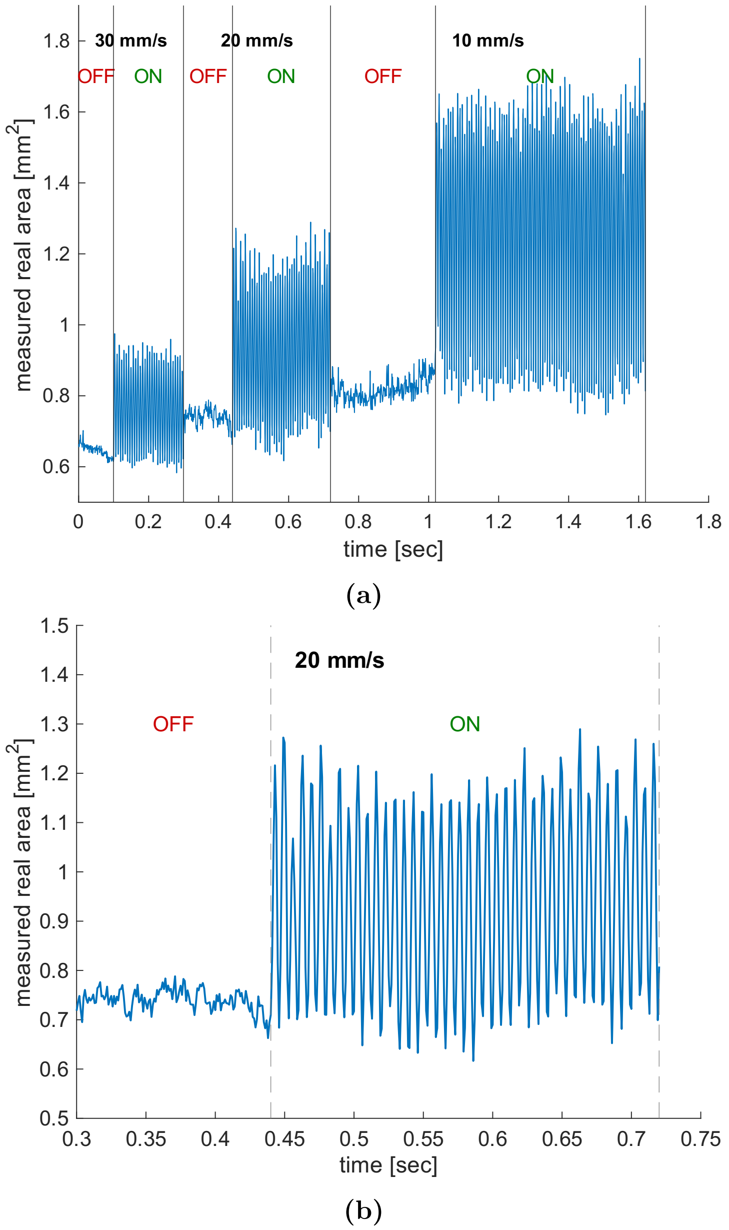


**Figure S5: Extracted real contact area data from fingerprint images of Participant-1 under an applied normal force of 1 N.** (a) Real area data when the camera trigger is on. (b) Real area measurement for 20 mm/s when the voltage is off and on

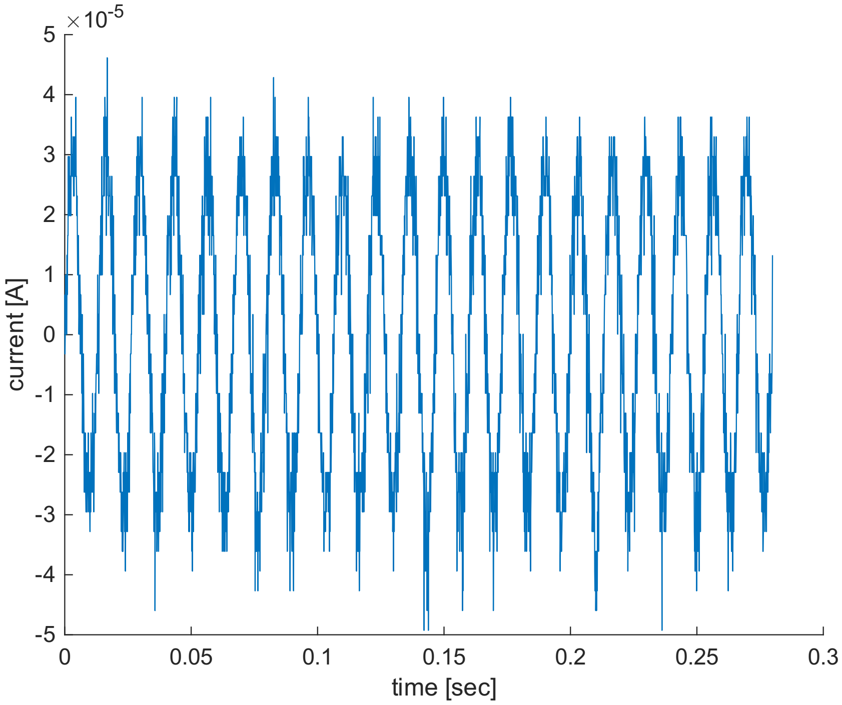


**Figure S6: Current measurement of Participant-1 under an applied normal force of 1 N.** Example of current data recorded when the camera and voltage-on trigger are on.

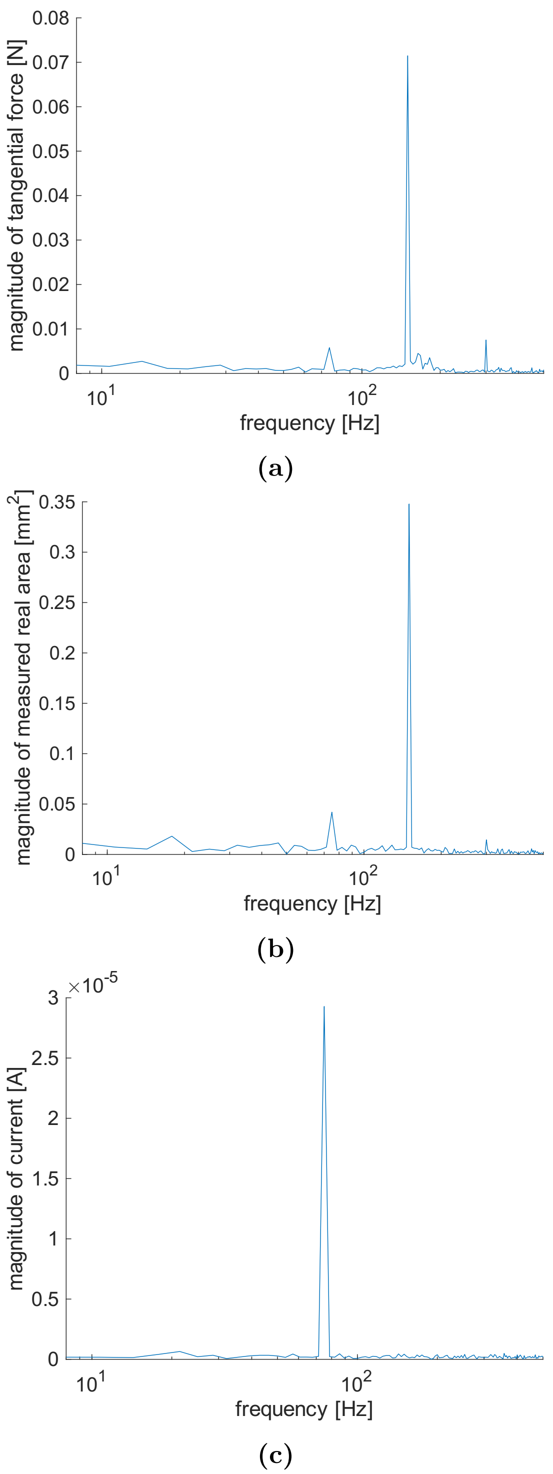


**Figure S7: Fourier transform of the recorded signals.** Fourier transform of (a) tangential force, (b) real area, and (c) current data during a trial with electrostatic actuation.

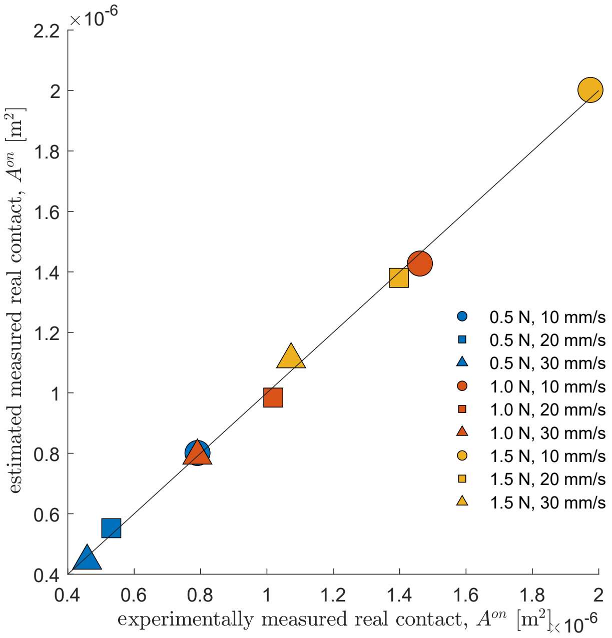


**(a)**

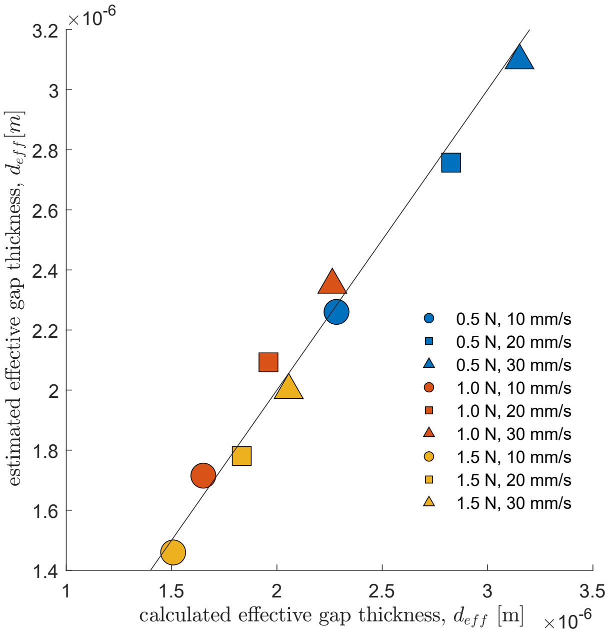


**(b)**

**Figure S8: Mean-level power-law parameterizations.** Measured-versus-estimated comparisons for the mean-level power-law parameterizations of measured real contact area $A^{\text{on}}$, effective gap thickness $d_{\text{eff}}$

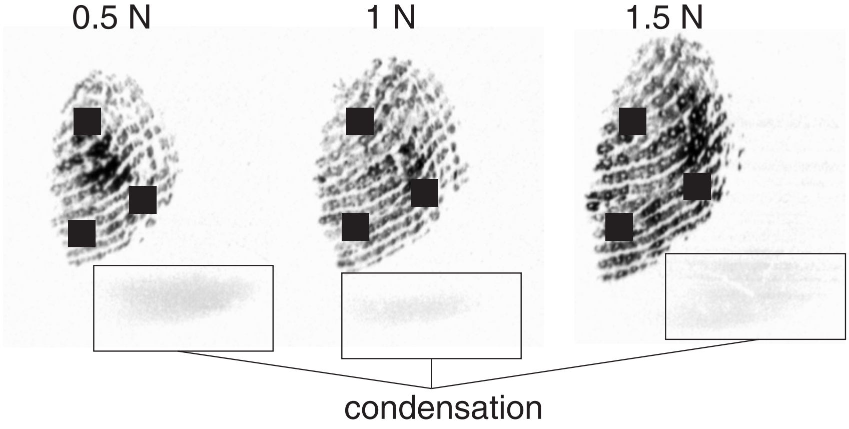


**Figure S9: Representative fingerprint images showing condensation, indicating the presence of moisture.**

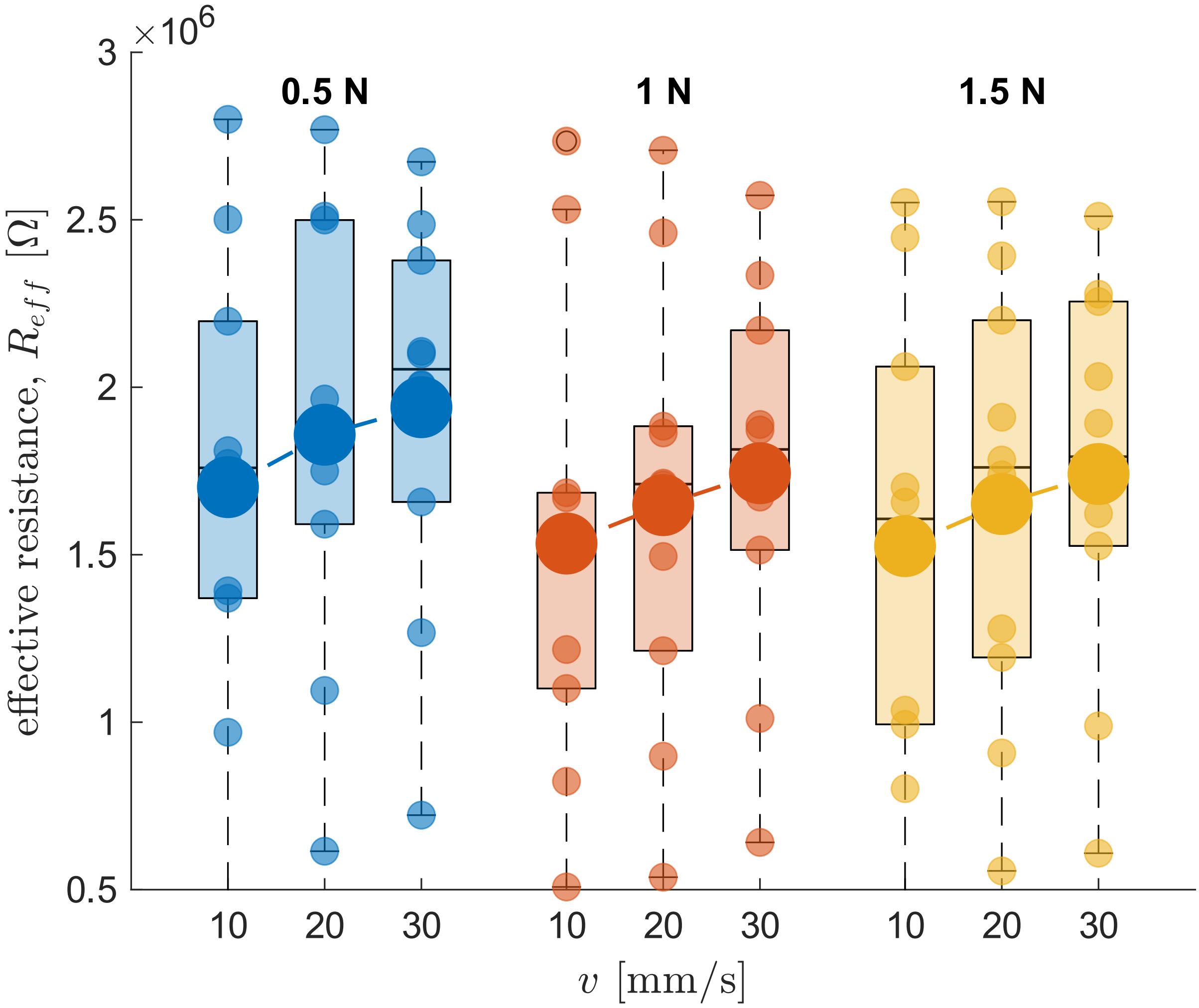


**Figure S10: Effective resistance parameters.** Result of $R_{eff}$ for different speeds and forces

### Supplementary Movies

**MovieS1** A representative video of fingerprint contact during a single trial at normal forces of 0.5 N, 1 N, and 1.5 N, with a sliding speed of 20 mm/s.

**MovieS2** A representative video of fingerprint contact during a single trial at normal forces of 10 mm/s, 20 mm/s, and 30 mm/s, with a sliding speed of 1 N.

**MovieS3** A representative video comparing moist and dry fingerprint contact during a single trial at a normal force of 1.5 N and a sliding speed of 10 mm/s.
